# Epigenetic age provides insight into tissue origin in endometriosis

**DOI:** 10.1101/815936

**Authors:** Katie Leap, Iveta Yotova, Steve Horvath, Julian A Martinez-Agosto

## Abstract

**Background:** Endometriosis is a common reproductive disease with a heterogeneous presentation. Classification attempts have thus far not offered insight into its cause or its symptoms. Endometriosis may result from the migration of shed endometrium to the peritoneal cavity. However, there are cases reported in girls without uteruses and men. While a non-retrograde menstruation origin of ectopic tissue is certain in these cases, we propose using DNA methylation age (DNAm age) to distinguish between retrograde and non-retrograde etiologies.

**Methods:** Using publicly available DNA methylation data and Horvath’s pan-tissue epigenetic clock, we compared DNAm age and epigenetic age acceleration (EAA) of ectopic lesions to eutopic endometrium of diseased and control endometrium. We examined EAA in cancer metastasis and teratomas to control for migration and developmental origin.

**Results:** Disease status does not change DNAm age of eutopic endometrium, but the effect of ectopic status was profound: −16.88 years (p = 4.82 × 10^−7^). There were no differences between EAA of primary/metastatic tumor pairs, suggesting that the observed effect is not due to migration. Immature or mature teratoma compartments decreased DNAm age by 9.44 and 7.40 years respectively, suggesting that developmental state correlates with DNAm age.

**Conclusions:** Ectopic endometriotic tissue exhibits decelerated DNAm age, similar to that observed in teratomas composed of multipotent tissue. The migration process does not change DNAm age and eutopic endometrium is concordant with chronological age regardless of disease status. We conclude that DNAm age of ectopic lesions can classify endometriosis into distinct subtypes that may be clinically relevant.

## Introduction

Endometriosis is a common reproductive disease that affects up to 10% of women defined by the presence of endometrium-like tissue in ectopic locations^1^, but the presence or size of ectopic endometriotic lesions does not predict fertility or the type or severity of pain experienced^2^. Attempts to classify the disease according to surgical, clinical or molecular features have thus far not offered any insight into what causes the disease or its symptoms^3^. We propose a classification system that uses DNA methylation age to predict tissue origin with the hope that this classification will enable better understanding of the disease and tailored treatment for affected women.

DNA methylation is an important part of gene regulation and is essential to development. During embryogenesis, the zygote undergoes dramatic DNA methylation changes, where many marks are erased and reestablished. DNA methylation continues to change throughout the lifespan of an individual in a predictable way such that the age of a person can be reliably predicted from the DNA methylation of a donated tissue^4^. One such method is Horvath’s pan tissue clock, which has as its input methylation values and as its output a predicted age in years^5^. Although DNA methylation patterns are variable between tissue types, the pan tissue clock applies to all nucleated cells.

Most tissues have roughly the same age according to the pan tissue clock with the exception of female breast tissue and the cerebellum ^5–7^. Furthermore, it predicts an embryological age, i.e. close to 0 years, for induced pluripotent stem cells, indicating that the biological process underlying the clock may be related to tissue status along the differentiation-senescence timeline^4^. We hypothesized that this property of the pan tissue clock (young age estimate for stem cells) might be beneficial in the context of establishing tissue origin, particularly for conditions like endometriosis in which this question remains a challenge.

Endometriosis is thought to be caused by the migration of shed endometrium to the peritoneal cavity through retrograde menstruation. Recent studies have demonstrated this disease mechanism by implicating mutations originating within the glands of the endometrium in the pathogenesis of endometriosis-associated ovarian cancer^8,9^. However, there are reported cases of endometriosis in girls who have not menstruated or who do not have uteruses, as well as in fetuses and in men^10–12^. While a non-retrograde menstruation origin of ectopic tissue is certain in these cases, we propose that the pan-tissue clock can distinguish between uterine and non-uterine etiologies in the commonly observed ambiguous cases. Here we demonstrate that ectopic endometriotic tissue exhibits a younger than expected (i.e. decelerated) DNAm age, similar to that observed in teratomas composed of multipotent tissue. The process of migration does not change DNAm age, and eutopic endometrium is concordant with chronological age regardless of disease status. We conclude that DNAm age of ectopic lesions can classify endometriosis into distinct subtypes that may be clinically relevant.

## Materials and Methods

### Data selection and study population

All data used in this analysis was publicly available from the Gene Expression Omnibus (GEO); accession numbers, sample sizes, and platform information can be found in Supplementary Table 1. Given how small the publicly available datasets of endometriosis tissues are, we had neither the power nor the variation necessary to study the entire methylome. Instead, we distilled the methylome into one biologically relevant metric and used larger datasetswith known cell migration or developmental properties to identify how the metric changes under these conditions. We chose datasets for the metastatic cancer analysis based on the following inclusion criteria: presence of tissue from both primary and metastatic tumors, availability of age information, and platform compatibility. To investigate tissues of a developmental origin, we used a dataset of intracranial germ cell tumors, which are classified into five different types: germinomas, embryonal carcinomas, teratomas, yolk sac tumors, and choriocarcinomas^13^. We focused on teratomas because of their developmental origin and their low likelihood of being malignant^14^. Patient characteristics of control and patient groups have been previously described{ref}.

### Horvath’s pan-tissue clock

We chose to use Horvath’s pan-tissue clock for the following reasons: i) it applies to all nucleated cells and tissues, ii) it lends itself to comparing the ages of different tissues^5,6^, iii) it leads to a very low age estimate in the case of stem cell types, and iv) it is validated in both 27k and 450k Illumina platforms^5^. Full details on DNA methylation age and epigenetic age acceleration calculated using Horvath’s pan-tissue clock are described elsewhere^5^ and the software for calculating these metrics is hosted online (https://horvath.genetics.ucla.edu/html/dnamage/). Briefly, DNA methylation age represents the predicted age of a tissue in years. If the clock is performing well, there will be a strong positive correlation between predicted age and actual age; we expect this correlation in healthy tissues, but relax the assumption for diseased tissues. Because age cannot be negative and is thus bounded at 0, there is more variability in predicted age as actual age increases. This means that the same difference between predicted and actual age in a child is more significant than in an older person. Therefore, we do not use raw differences to represent age acceleration (or deceleration), and instead use the residual from the model predicting DNAm age from actual age, which we call epigenetic age acceleration and can be positive or negative. The sensitivity of the clock is 3.6 years as determined by the median absolute difference between DNAm age and chronological age in the testing data used to develop the method.

### Statistical analyses

Before conducting any analyses, we confirmed correlation of DNAm age with chronological age using Pearson’s correlation coefficient as well as graphically (Figure 1A). We used epigenetic age acceleration (EAA) initially to compare different tissue types graphically with boxplots. However, we chose to use age as a covariate and predict DNAm age using ordinary least squares regression because age was missing from some of our samples and EAA cannot be calculated if age is missing. We fit three different models: a) an endometriosis model with the following covariates: age, diagnosis of endometriosis (binary), menstrual phase (binary: 0 for secretory, 1 for proliferative), ectopic status (binary), and stromal status (binary); b) a metastatic cancer model with the following covariates: age and metastatic status via three binary indicator variables: primary tumor with metastases, primary tumor without metastases, and metastatic tumor; and c) a teratoma model with the following covariates: age, tumor content of the sample (percentage) and non-mutually exclusive binary indicator variables for embryonic carcinoma, germinoma, immature teratoma, mature teratoma, seminoma and yolk sac tumor. Age estimates for each model can be seen in Figure 1B. The Akaike Information Criterion (AIC) was used to select the best model. Significance for each coefficient was determined using a t-test and confidence intervals were bootstrapped. Paired analyses of DNAm age were conducted using paired t-tests. Effect sizes smaller than 3.6 years were not considered significant because of the sensitivity of the epigenetic clock. We did not correct for multiple testing because we used one model for each analysis, but we have reported raw p-values.

**Figure 1.**
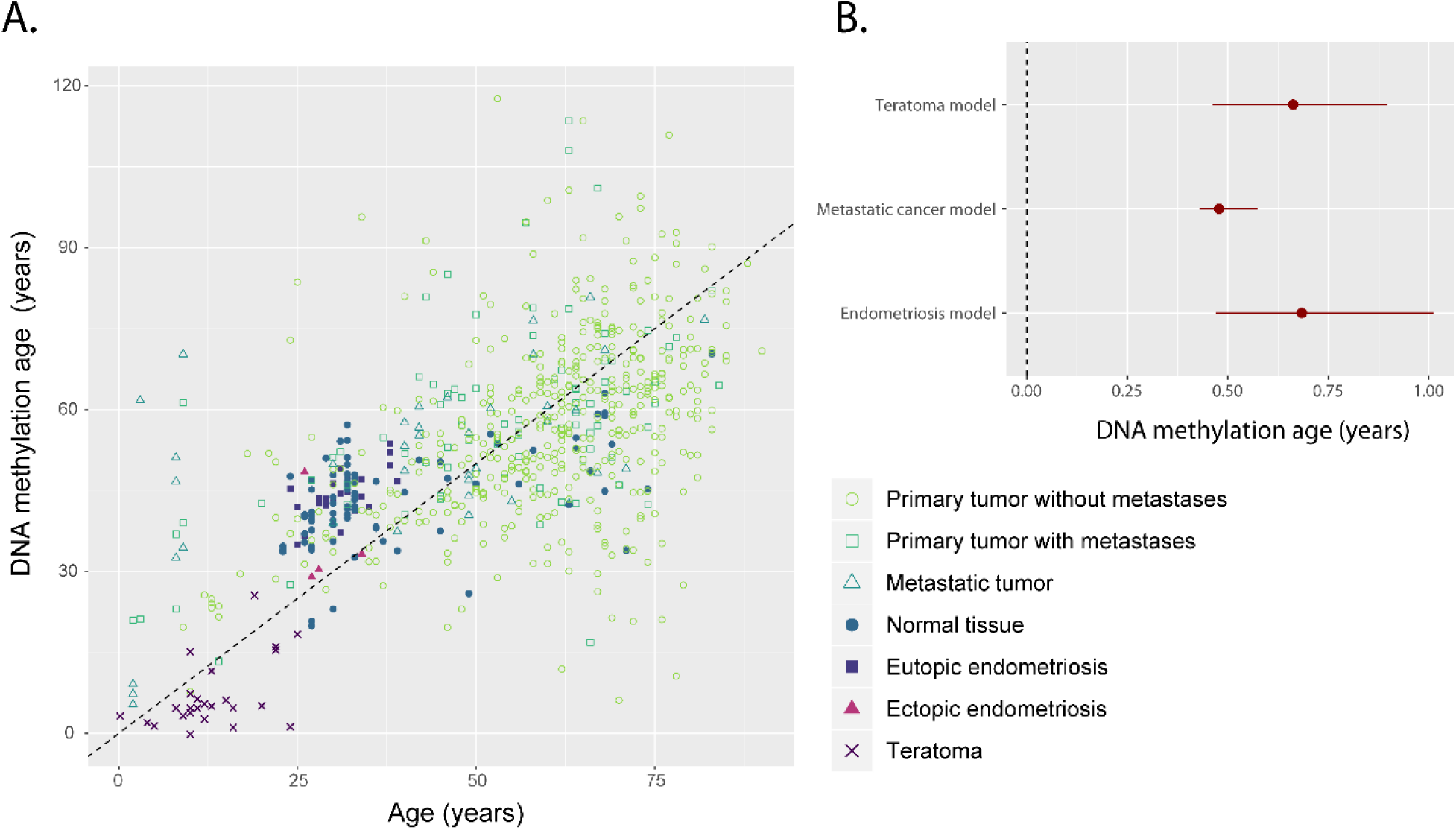
A) Correlation between chronological age and predicted DNA methylation age. Dashed line indicates the identity line where perfectly correlated points would be found. Open shapes are cancerous tissues; filled shapes are non-cancerous. B) Age parameters estimated from each of the three multivariate models; point estimate and bootstrapped 95% confidence interval are shown. A value of 0 would indicate that age does not affect DNAm age; a value of 1 would indicate that for every 1 year increase in age, DNAm age increases by 1 year.

### Missing data

Out of 685 samples in the metastatic cancer dataset, 88 were missing metastatic status, most frequently because stage of the cancer was available but no information about the metastatic status of the particular tumor sample was available. The difference in DNAm age between the tissues with known metastatic status and unknown metastatic status was 2.4 years (p = 0.27). As the DNAm age was not significantly different and was within the margin of error of the clock (3.6 years), we considered these to be missing at random and excluded them from our analyses.

## Results

### Endometriosis does not affect DNA methylation age of eutopic endometrium

We first tested whether the DNAm age of eutopic endometrium of women with and without endometriosis is concordant with chronological age using Pearson’s r and found a positive correlation of r = 0.6 (p = 7 × 10^−6^), which is consistent with previous findings in uterine endometrium^5^. A recent study found a correlation of 0.8 between DNAm age and chronological age in uterine endometrium when all of the samples came from the same menstrual time point (LH+7, or 7 days after the surge in luteinizing hormone)^15^, but our correlation was unchanged if we stratified by menstrual phase (LH+8), or by endometriosis status. We next tested the effect of menstrual phase on DNAm age because menstrual phase is known to affect DNA methylation^16^. Each of the healthy controls donated two samples, one during the early secretory phase and the second during mid-secretory phase of the same menstrual period^17^. These two time periods are just before and just after the differentiation of the stromal cells to decidual cells at the beginning of the receptive phase of the endometrium^18^. If the clock is tracking differentiation, we might expect an older age after this change. The average difference in DNAm age between these paired samples was 2 years (p = 0.07), which is within the sensitivity of the clock, and the correlation between the two DNAm ages was 0.7 (p = 0.001), a stronger correlation than with chronological age (Figure 2). This implies that menstrual phase has a limited effect on DNAm age. Finally, we tested the repeatability of DNAm age: the mid-secretory phase samples each had a technical replicate and the correlation between the DNAm age of these paired replicates was incredibly strong at r = 0.99 (p = 4 × 10^−15^). Given that DNAm age is correlated with age in the eutopic endometrium, menstrual phase has a small effect, and the predicted age is repeatable, we tested whether endometriosis would affect DNAm age of the eutopic endometrium. In our full linear regression model, we predicted DNAm age given chronological age, ectopic status, endometriosis status, menstrual phase, and stromal cell status. The effect of disease status was not significant (0.98 years, p = 0.344) indicating that disease status does not change DNAm age in the eutopic endometrium (Figure 2A).

**Figure 2.**
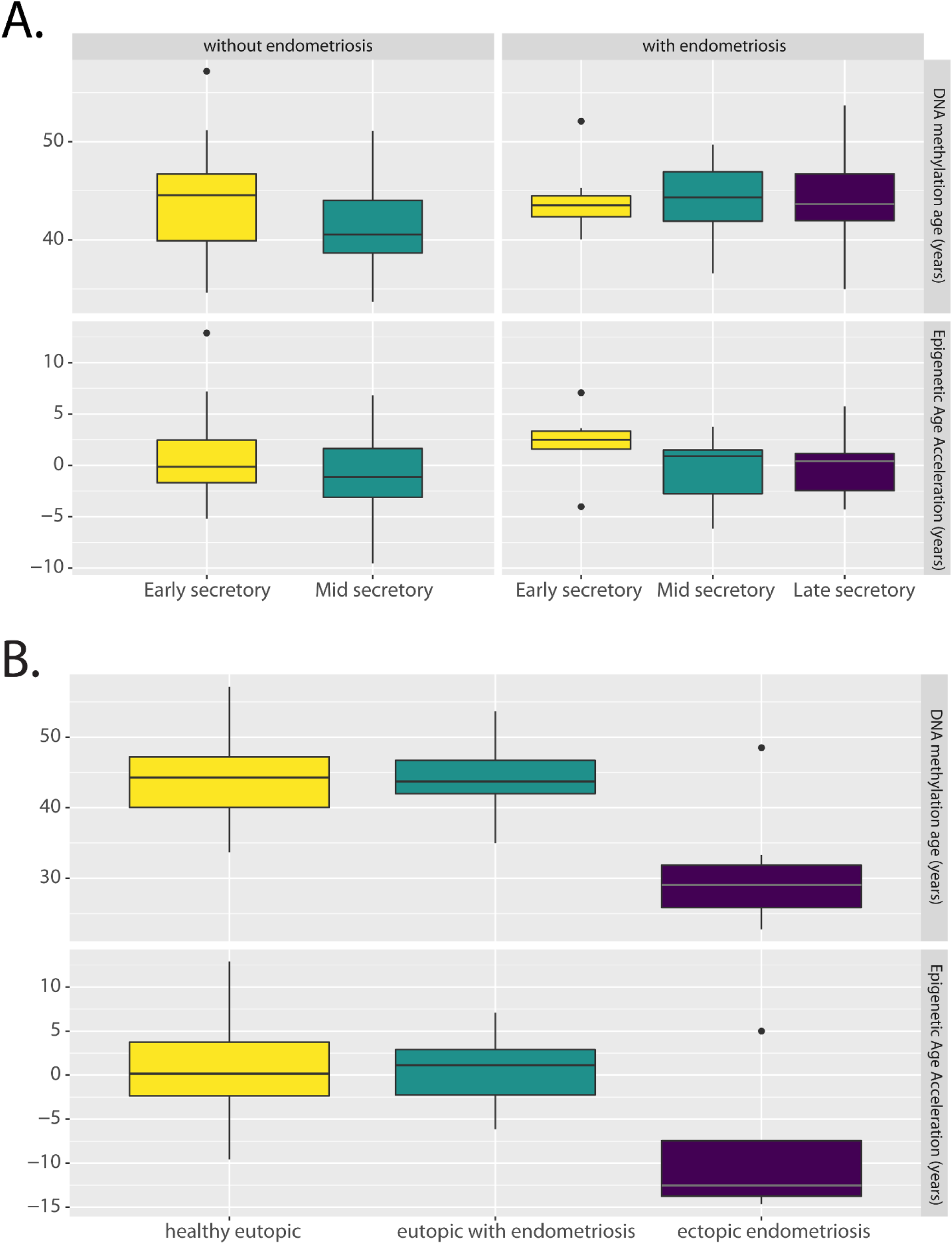
A) Menstrual phase and disease status do not have a significant effect on DNA methylation age (top) or Epigenetic Age Acceleration (bottom). All samples are of eutopic tissue, coming from 26 women with endometriosis (early secretory: n = 7, mid secretory: n = 10, late secretory: n = 9) and 17 women without endometriosis who each donated samples at early- and mid-secretory phases of the same menstrual cycle. B) Ectopic endometriotic lesions are significantly younger than eutopic samples as seen in the boxplot; midline is the median, box edges represent the interquartile range. Healthy eutopic: n = 42; eutopic with endometriosis: n = 29; ectopic endometriosis: n = 7.

### Ectopic endometriotic tissue is epigenetically younger than eutopic endometrium

We next compared the DNAm age of ectopic endometriotic lesions with that of eutopic tissue. Ectopic tissue was substantially younger than expected, exhibiting a profound age deceleration: −16.88 years (p = 4.82 × 10^−7^, Figure 2B). As ectopic lesions are often located on the ovary, as were all of the samples used in this analysis, we could not completely exclude adjacent tissue contamination, which might consist of ovarian epithelium, stroma, or germ cells. However, it is unlikely that the effect of ectopic status on DNAm age reflects adjacent ovarian tissue contamination for the following reasons. First, by construction, the pan-tissue clock is quite robust to differences in cell composition^5^. Second, the ectopic endometriosis data came from primary stromal cells isolated from surgical tissue samples thereby representing a pure population of cells, grown ex vivo for not more than three passages^19^. Third, we trained a logistic regression classifier on the full methylome to predict whether a tissue was ovarian or not and determined that the ectopic lesions were not likely to be ovarian. Overall, it is unlikely that the substantially younger DNA methylation age of ectopic tissue reflects artifactual contamination with ovarian tissue.

### Stromal cell identity does not explain age deceleration

Because all of the ectopic samples consisted of cultured stromal cells, we considered whether cultured stromal cells might exhibit age deceleration. Mesenchymal stromal cells derived from human bone marrow have previously been shown to have a DNAm age that is correlated with chronological age and therefore lacking any age deceleration. When these cultured stromal cells were reprogrammed to become induced pluripotent stem cells (iPSCs), their DNAm age decreased to zero, but differentiation back into mesenchymal stromal cells did not increase their DNAm age^20^. This supports our finding in the endometrium that differentiation does not advance Horvath’s pan-tissue clock and does not give us reason to suspect that stromal cell identity could be driving ectopic epigenetic age deceleration. We examined eutopic stromal cells to see if they exhibit any epigenetic age acceleration or deceleration as compared to the whole tissue (Figure 3A). Our results found that if we were to hold all other variables constant, stromal cells would be on average 6.57 years older than the whole tissue, with a 95% confidence interval estimate of 0.68-11.62, p = 0.0138. This age acceleration could be due to the effect of passaging, which has previously been shown to advance DNAm age^21^. Incorporating stromal status in the full model as a covariate did not change the effect size of ectopic status, indicating that while stromal samples may be older on average than whole tissue, this effect is independent of the age deceleration seen with ectopic status (Figure 3B).

**Figure 3.**
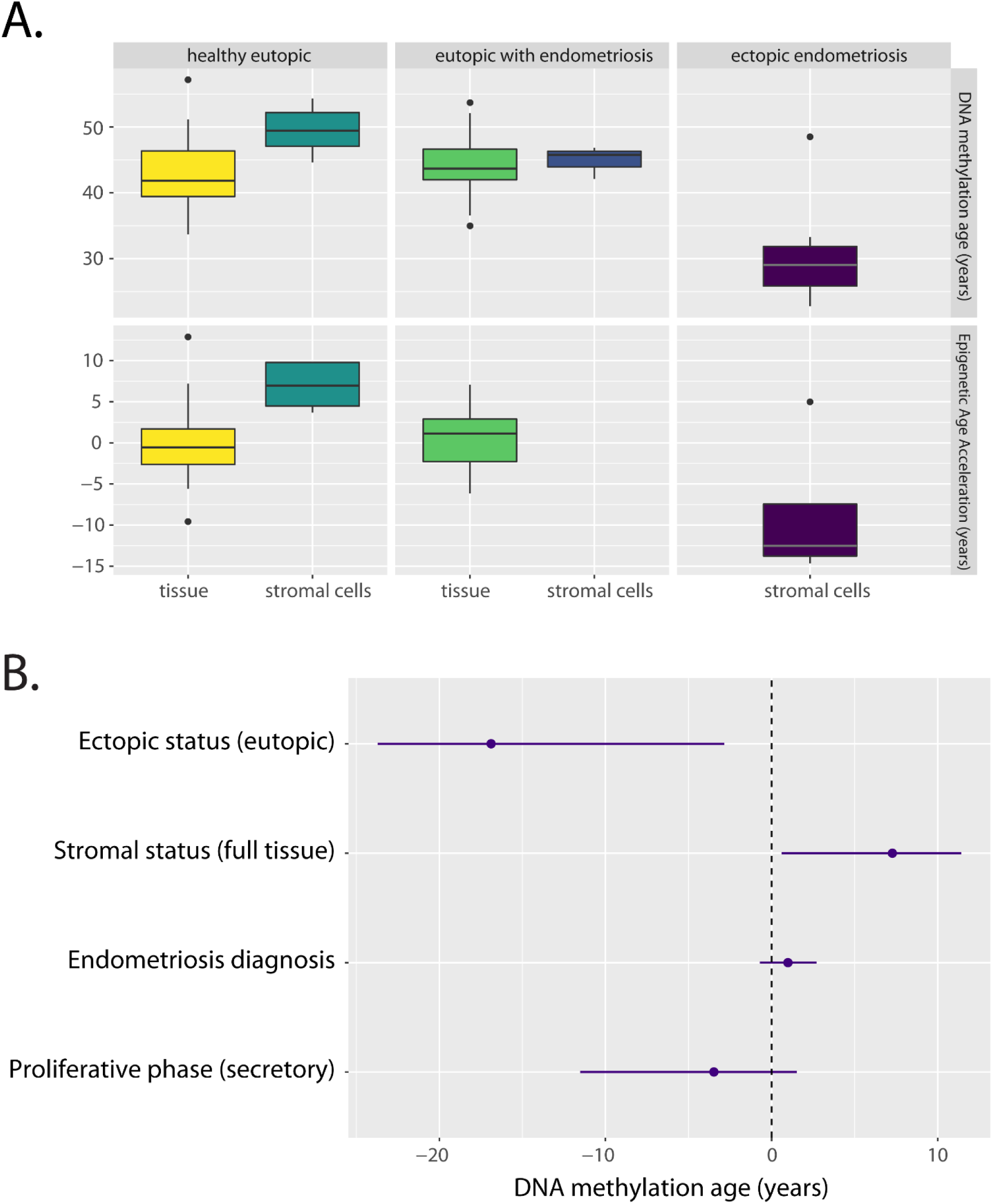
A) DNA methylation age and Epigenetic Age Acceleration (EAA) of whole tissue as compared to isolated stromal cell populations. The age acceleration observed in stromal cells may be due to the effect of passaging. Healthy tissue: n = 34; Healthy stromal cells: n = 8; Eutopic tissue (endometriosis): n = 26; Eutopic stromal cells (endometriosis): n = 3; Ectopic stromal cells: n = 7. B) Values of the coefficients estimated from the multivariate linear regression model; the point estimates are shown as points while the lines represent a bootstrapped 95% confidence interval. If the line crosses 0, the effect is not significant at 95% confidence. Age coefficient can be found in Figure 1B.

### Cell migration does not affect DNA methylation age

To examine the effect of migration processes on DNAm age, we analyzed datasets of metastatic cancers as metastasis is a prototypical example of (malignant) tissue migration. Because it is known that cancer can dysregulate DNAm age, we confirmed the performance of Horvath’s pan-tissue clock through the correlation between chronological age and DNAm age in the normal tissue samples: r = 0.85 (p = 3 × 10^−10^). As seen in Figure 4A, cancer samples were more variable than normal tissues and exhibited age acceleration on average. However, the epigenetic age acceleration of metastatic tumor samples did not significantly differ from that of primary tumor samples on aggregate or in matched paired analyses, as can be seen in Figure 4B by comparing the parameter estimates of primary tumors with metastases to metastatic tumors. This suggests that while the process by which a tissue becomes cancerous and invasive may accelerate its DNAm age, migration alone is not enough to affect DNAm age. These findings exclude the cell migration process or ectopic location as a contributor to DNAm age deceleration in ectopic endometriotic tissues.

**Figure 4.**
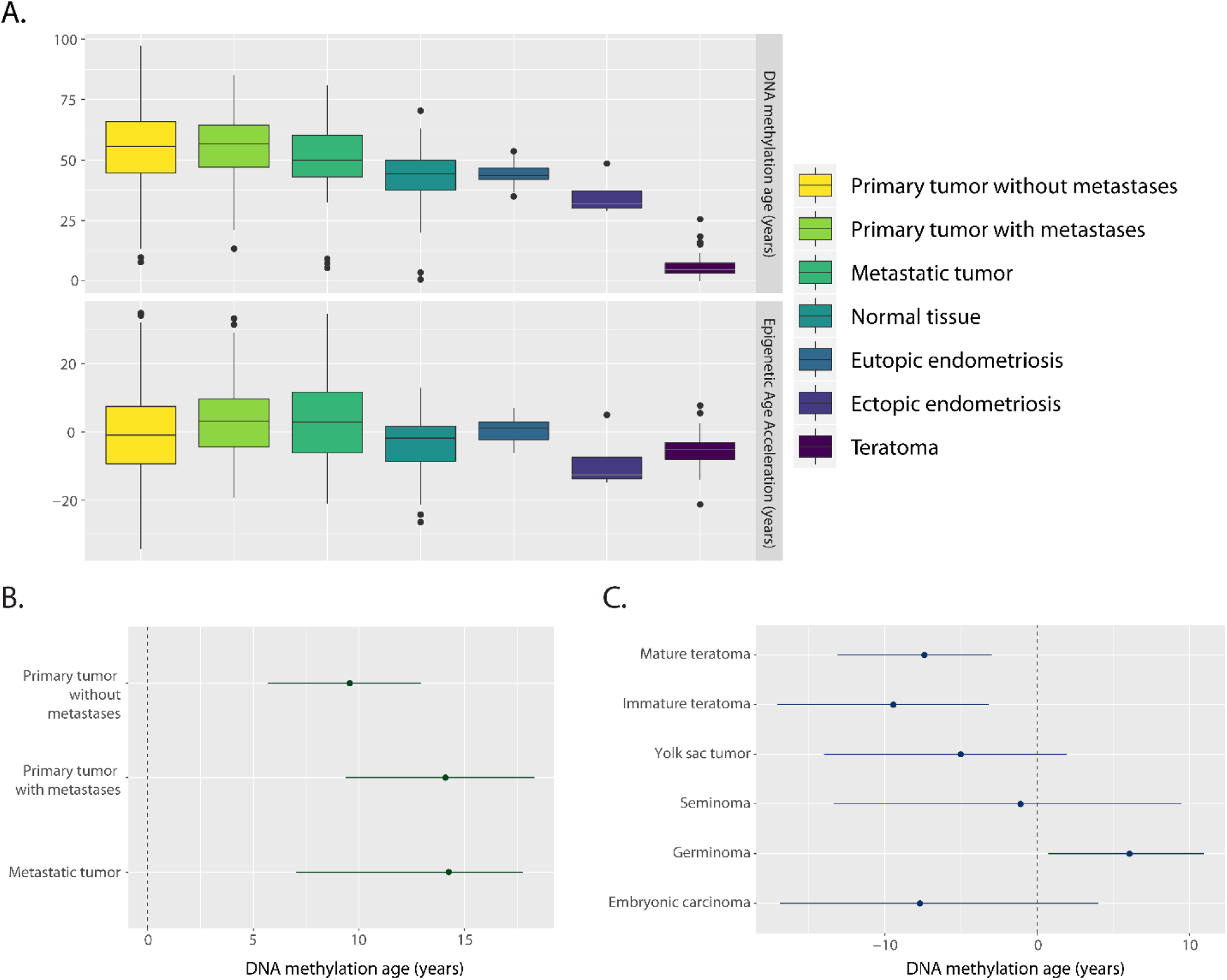
A) Predicted DNA methylation age and epigenetic age acceleration by tissue type. Ectopic endometriotic tissue shows age deceleration comparable to that seen in teratomas. Primary tumor without metastases: n = 534; Primary tumor with metastases: n = 82; Metastatic tumor: n = 38; Normal: n = 81 (8); Eutopic endometriosis: n = 29 (3); Ectopic endometriosis: n = 7 (3); Teratoma: n = 25. Since EAA is dependent on knowledge of donor age, the samples for which the information about the age was absent were excluded from EAA analyses. These samples are given in parentheses. Limits of EAA plot exclude 23 outliers: 17 in primary tumors without metastases (3%), 5 in primary tumors with metastases (6%) and 1 in metastatic tumors (3%). B & C) Values of the coefficients estimated from multivariate linear regression models; the point estimates are shown as points while the lines represent a bootstrapped 95% confidence interval. If the line crosses 0, the effect is not significant at 95% confidence. Age coefficients can be found in Figure 1B.

### Endometriotic stromal cells share a similar DNA methylation age pattern with multipotent tissues

Since cell migration, such as retrograde menstruation, does not explain the substantial DNAm age deceleration of ectopic tissue, we explored the possibility that ectopic tissue had been deposited during development and might share a similar DNAm age to tissues of developmental origin. Teratomas are benign growths of multipotent tissue that originate from developmental abnormalities, are always ectopically located, and are composed of undifferentiated cell types. We hypothesized that teratomas would exhibit DNAm age deceleration, but not as much as is observed in iPSCs or progenitor cells. The linear regression model we used to predict DNAm age of germ cell tumors given chronological age, tumor content, and indicator variables for tumor types was able to explain 75% of the variance in DNAm age. The presence of immature teratomas or mature teratomas decreased the predicted DNAm age by 9.44 and 7.40 years respectively (p = 0.0016 and p = 0.0074). While the teratomas were predicted to have a very young DNAm age (Figure 4A), this is explained by the fact that most of the samples came from young children (range 2 months-25 years, Figure 1A). When adjusted for age with EAA, the difference is not as drastic (Figure 4A) and the teratomas are more similar to ectopic endometriotic tissue.

Interestingly, embryonic and placental germ cell tumors (embryonal carcinoma and yolk sac tumors) did not exhibit the age deceleration seen in mature and immature teratomas (Figure 4C). Furthermore, none of the teratoma samples exhibited a DNAm of zero, as would be expected in fetal cells or iPSCs^5^. Taken together, this implies that tissue of a developmental origin exhibits epigenetic age deceleration that is not as extreme as is seen in fetal tissues or iPSCs, which is the pattern seen in ectopic tissue in endometriosis, and thus ectopic tissue in endometriosis may have a developmental origin.

## Discussion

We show that epigenetic age estimators (e.g. the pan tissue epigenetic clock by Horvath) lend themselves to addressing vexing problems surrounding the etiology of endometriosis. Using human tissue samples, we demonstrate a) that DNAm age strongly correlates with chronological age in the endometrium, b) that ectopic tissue is substantially younger than eutopic tissue according to the epigenetic clock, c) that tissue migration through metastasis does not change DNAm age, and d) that ectopic endometriotic lesions and teratomas share age deceleration that is not as young as pluripotent embryonic or induced pluripotent cells according to the epigenetic clock.

Endometriosis is primarily a young woman’s disease; symptoms often begin during puberty and the vast majority of cases resolve with menopause^22^. Senescence is a hallmark of aging, but it is also important for development and tissue homeostasis^23^. Endometriosis can be thought of as a problem of senescence because it is ultimately a problem of growth: endometriotic tissue should not grow and proliferate in ectopic sites, regardless of the exact process by which this growth occurs or is prevented in a non-diseased state. Therefore, it is possible that the natural process of aging causes senescence in ectopic tissues that had previously been resistant to apoptosis and necrosis. Using compounds to change DNAm age of ectopic lesions to one more consistent with a differentiated cell type may represent a therapeutic approach, akin to the success of differentiation therapy in leukemias^24^.

Variability between lesions within person has been well-documented^16^ and future research should investigate variability of DNAm age between lesions within an individual. While all of the lesions studied here were endometriomas, peritoneal disease presents with different colored lesions that are thought to represent a continuum from younger to older disease^25^, a hypothesis that could be explored using DNAm age.

Epigenetic clocks have potential utility as a prognostic indicator. Tissue of an embryonic origin may have a different clinical course than tissue resulting from retrograde menstruation and may necessitate different surgical or medical interventions. In order to validate this hypothesis, samples from women with known retrograde flow, such as that resulting from imperforate hymen, should be compared to those from affected men or women without a uterus. These disparate extremes may shed light into the etiologies of endometriosis and better methods of diagnostics, classification, and treatment.

## Acknowledgements

The authors declare no conflict of interest. This work was supported by a NIH-NCI grant T32LM012424, Biomedical Big Data.

---

**Supplemental Table 1:**

| **GEO Accession** | **Tissue Type** | **Sample Size** | **Platform** | **Analysis** |
| --- | --- | --- | --- | --- |
| GSE34355 | medulloblastoma | 15 | GPL8490 | metastatic cancer |
| GSE73832 | small intestine cancer | 97 | GPL13534 |  |
| GSE62231 | paraganglioma | 123 | GPL8490 |  |
| GSE77269 | hepatocellular carcinoma | 57 | GPL13534 |  |
| GSE53051 | multiple cancers | 220 | GPL13534 |  |
| GSE39279 | non-small lung cancer | 515 | GPL13534 |  |
| GSE93589 | endometrial cancer | 9 | GPL13534 |  |
| GSE81224 | high grade serous ovarian cancer | 20 | GPL13534 | ovarian classifier |
| GSE43265 | ovarian cancer | 31 | GPL8490 |  |
| GSE26989 | ovarian cancer | 51 | GPL8490 |  |
| GSE73948 | endometrium (endometriosis) | 24 | GPL13534 | endometriosis |
| GSE73949 | endometrium | 17 | GPL13534 |  |
| GSE73949 | endometrium | 34 | GPL13534 |  |
| GSE87621 | stromal cells | 9 | GPL13534 |  |
| GSE47359 | stromal cells | 9 | GPL8490 |  |
| GSE70783 | intracranial germ cell tumors | 77 | GPL13534 | teratomas |

GPL8490: Illumina HumanMethylation27 BeadChip
GPL13534: Illumina HumanMethylation450 BeadChip

